# The _329_HHK_331_ motif is essential for Alzheimer’s disease tau filament fold

**DOI:** 10.64898/2026.06.29.735439

**Authors:** Yuta Sato, Masato Kawasaki, Toshio Moriya, Miki Senda, Masami Masuda-Suzukake, Kanae Ando, Shin-ichi Hisanaga, Masato Hasegawa, Toshiya Senda, Takashi Nonaka

## Abstract

Cryo-electron microscopy (cryo-EM) has revealed disease-specific tau filament folds, yet the local sequence elements that determine them remain poorly understood. Here we focused on the _329_HHK_331_ motif near an inter-protofilament interface in Alzheimer’s disease (AD)-type tau filaments, and analyzed recombinant dGAE filaments of wild-type (WT) and mutants in this motif. All mutants formed amyloid-like filaments *in vitro*, but their morphologies differed. In cultured cells, WT filaments efficiently seeded WT tau aggregation. Filaments with three-residue changes (deletion or Ala substitution) showed almost no seeding activity, whereas two-residue deletions retained partial activity. Cryo-EM showed that WT dGAE filaments form a quadruple helical filament of two protofilament dimers. Each dimer comprises two C-shaped protofilaments, centered on a _333_GGG_335_-mediated inter-protofilament interaction and supported by flanking _329_HHK_331_-_336_QVE_338_ contacts. Three-residue alterations abolished interactions with the QVE motif at the protofilament interface, thereby displacing _333_GGG_335_ and forming non-C-shaped protofilament structures that are intrinsically poor templates for tau seeding. By contrast, two-residue deletions maintained the C-shaped protofilament structure because the remaining residue formed alternative inter-protofilament interactions. These findings suggest that the _329_HHK_331_ region is a key determinant of the AD-like C-shaped protofilament fold and link this motif to tau seeding, providing insight into disease-specific tau filament formation.

## Introduction

Alzheimer’s disease (AD) is a neurodegenerative disorder pathologically characterized by extracellular accumulation of aggregated amyloid β (Aβ) and intracellular accumulation of phosphorylated tau proteins^1^. These pathological aggregates form β-sheet-rich amyloid filaments that can act as seeds to recruit soluble proteins, thereby promoting filament elongation and the propagation of the pathology^2^. Cryo-electron microscopy (cryo-EM) studies have revealed that tau filaments isolated from the brains of patients with AD and other tauopathies adopt disease-specific folds^3, 4, 5, 6, 7^. These findings suggest that tau filament structure is closely associated with the molecular pathology of each tauopathy. However, how different folds are formed depending on the type of tauopathy remains unclear.

Recently, recombinant tau fragments comprising residues 297–391 (numbered according to the longest 441-residue tau isoform and referred to as dGAE tau, Fig. 1A) were reported to reproduce the disease-specific folds observed in patients with AD and chronic traumatic encephalopathy (CTE)^8^. In tau, the most widely recognized aggregation-promoting motif is PHF6 (_306_VQIVYK_311_), which serves as a critical nucleation site for filament formation^9^. Time-resolved cryo-EM analyses have revealed that during early stages of tau filament maturation in AD and CTE, a 15-residue segment including PHF6 forms filaments with a cross-α structure, suggesting a crucial role for PHF6 in initiating tau aggregation^10^. Several other sequences have been proposed as aggregation motifs, such as PAM4 (_350_VQSKIGSLDNITH_362_), _326_GNIHHK_331_, and _344_LDFKDR ^11^. PAM4 is regarded as a novel amyloidogenic motif and a major factor contributing to tau polymorphism, while the other sequences have only been studied for their general amyloid-forming potential and require further detailed investigation. Furthermore, the specific amino acid residues responsible for inducing disease-specific folding in tau filaments remain poorly understood.

**Figure 1.**
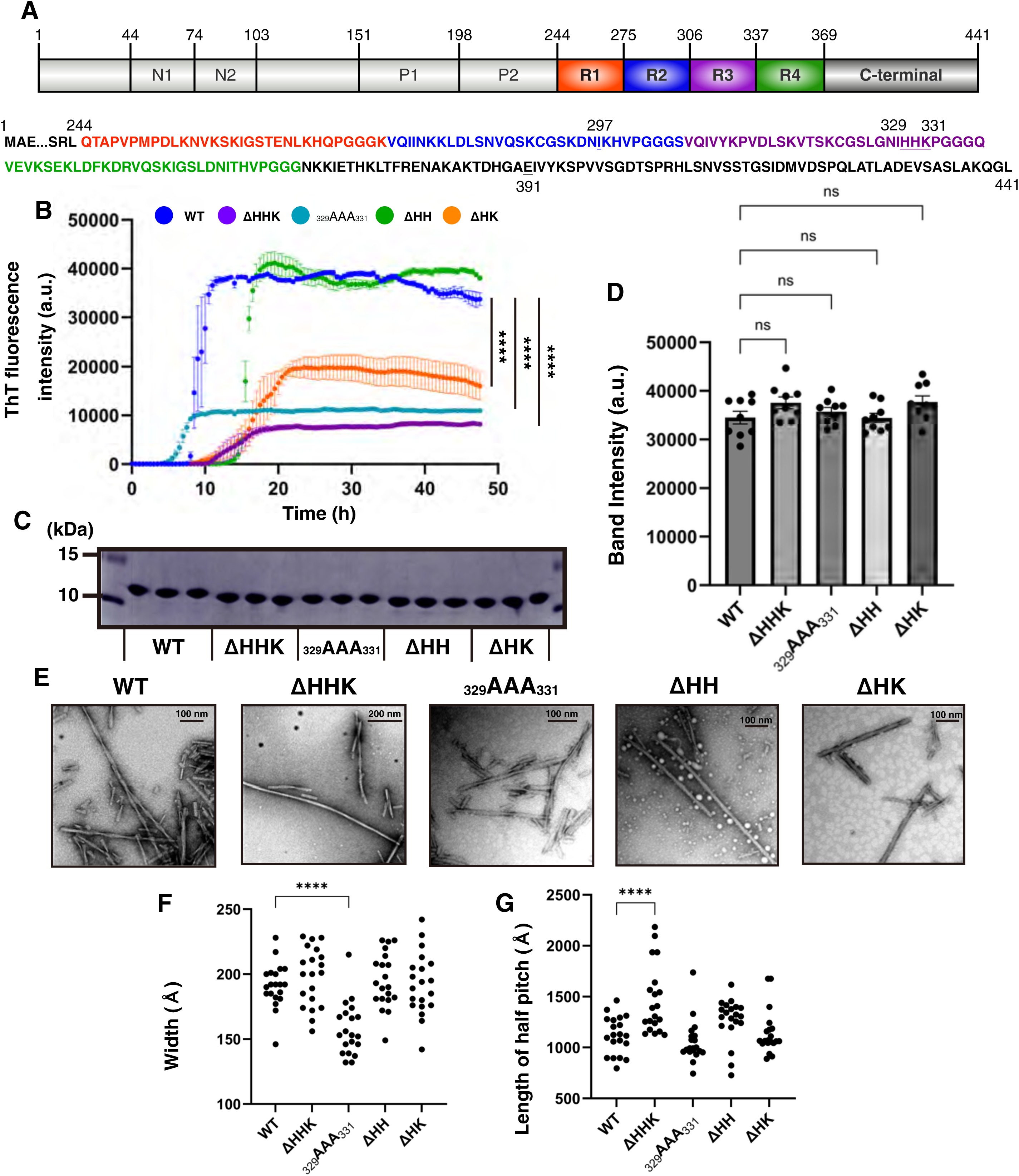
*In vitro* assembly and morphological characterization of wild-type and mutant tau filaments. (A) Schematic representation and amino acid sequence of the longest human 4R2N tau isoform. Top: domain organization with N1 (44–73), N2 (74–102), P1 (151–197), and P2 (198–243) in gray; R1–R4 (244–368) in red, blue, purple, and green, respectively; and the C-terminal region (369–441) in dark gray. Bottom: amino acid sequence for residues 244–441. The dGAE fragment (297–391) and the _329_HHK_331_ motif are underlined. (B) Fibrillization kinetics of recombinant tau proteins monitored by Thioflavin T (ThT). Data for WT (blue), ΔHHK (purple), _329_AAA_331_ (cyan), ΔHH (green), and ΔHK (orange) are plotted as mean ± SEM (N = 3). Final ThT fluorescence intensities were compared with WT using one-way ANOVA followed by Dunnett’s multiple comparisons test. ****, P < 0.0001. (C, D) SDS-PAGE analysis of dGAE tau filaments at ThT assay endpoint (C) and quantification of band intensity (D). Data represent mean ± SEM (N = 9; One-way ANOVA with Dunnett’s test, each tau mutant compared to WT). ns, not significant. (E-G) Negative-stain electron microscopy images of dGAE tau filaments (E) and quantification of filament width (F) and half pitch length (G). Each tau mutant was compared with WT (One-way ANOVA with Dunnett’s test; N = 20; mean ± SEM). ****, P < 0.0001. Scale bar, 100 or 200 nm.

In this study, we focused on the _329_HHK_331_ motif, which lies within an inter-protofilament interaction region of AD paired helical filaments (PHF)^5^ and immediately precedes the _332_PGGG_335_ motif, a region implicated in conformational rearrangements during AD tau filament maturation^10^. The _329_HHK_331_ motif contains clustered basic residues and has also been implicated in the binding of negatively charged aggregation-promoting factors, such as heparin^12^. We therefore hypothesized that this motif may play a specific structural role in the formation of AD-like tau filaments.

To test this hypothesis, we generated a series of recombinant dGAE tau mutants in which the _329_HHK_331_ motif was deleted or substituted, and examined the *in vitro* aggregation propensity, cellular seeding activity, and cryo-EM structures. By comparing WT dGAE tau filaments with mutants carrying three-residue alterations or two-residue deletions in the _329_HHK_331_ motif, we aimed to determine whether this region functions as a local sequence determinant of the AD-like C-shaped tau protofilament fold.

## Results

### Characterization of mutant dGAE tau filaments

To examine the role of the _329_HHK_331_ motif in tau filament formation, we generated recombinant dGAE tau mutants targeting this motif. These comprised ΔHHK (all three _329_HHK_331_ residues deleted), _329_AAA_331_ (the motif replaced with three alanines), and the two-residue deletion mutants ΔHH (His329 and His330 deleted) and ΔHK (His330 and Lys331 deleted) (Fig. 1A, Table 1).

**Table 1.** *In vitro* assembly conditions for each tau filament.

|  | Construct | Buffer | Shaking speed (rpm) | Shaking Method | Time (h) |
| --- | --- | --- | --- | --- | --- |
| dGAE tau WT | 297-391 | 10 mM Phosphate buffer pH 7.4, 10 mM DTT, 200 mM MgCl<sub>2</sub> | 200 | Orbital | 48 |
| dGAE tau <sub>329</sub>AAA<sub>331</sub> | 297-391, Alanine substitution of <sub>329</sub>HHK<sub>330</sub> | 10 mM Phosphate buffer pH 7.4, 10 mM DTT, 200 mM MgCl<sub>2</sub> | 200 | Orbital | 48 |
| dGAE tau ΔHHK | 297-391, deletion of <sub>329</sub>HHK<sub>331</sub> | 10 mM Phosphate buffer pH 7.4, 10 mM DTT, 200 mM MgCl<sub>2</sub> | 200 | Orbital | 48 |
| dGAE tau ΔHH | 297-391, deletion of <sub>329</sub>HH<sub>330</sub> | 10 mM Phosphate buffer pH 7.4, 10 mM DTT, 200 mM MgCl<sub>2</sub> | 200 | Orbital | 48 |
| dGAE tau ΔHK | 297-391, deletion of <sub>330</sub>HK<sub>331</sub> | 10 mM Phosphate buffer pH 7.4, 10 mM DTT, 200 mM MgCl<sub>2</sub> | 200 | Orbital | 48 |
| dGAE tau <sub>347</sub>AAA<sub>349</sub> | 297-391, Alanine substitution of <sub>347</sub>KDR<sub>349</sub> | 10 mM Phosphate buffer pH 7.4, 10 mM DTT, 200 mM MgCl<sub>2</sub> | 200 | Orbital | 48 |

WT dGAE tau has been reported to form PHF-like structures when incubated with MgCl_2_ under agitation at 37°C^8^. We therefore examined whether mutant dGAE tau proteins could form amyloid-like filaments under the same conditions. Filament formation was monitored by Thioflavin T (ThT) fluorescence. All mutant proteins showed time-dependent increases in ThT fluorescence, indicating that they retained the ability to form β-sheet-rich amyloid-like assemblies. However, ThT fluorescence intensities at plateau levels differed among mutants (Fig. 1B). The amount of aggregated tau, quantified by SDS–PAGE, was comparable across samples (Fig. 1C, D). Negative-stain electron microscopy further confirmed the formation of filamentous assemblies by all mutants, but revealed differences in filament morphology, width, and half-pitch length depending on the mutation (Fig. 1E–G).

Together, these results indicate that mutations in the _329_HHK_331_ motif do not abolish dGAE tau filament formation, but alter the biophysical and morphological properties of the resulting filaments. These findings suggest that WT and mutant dGAE tau filaments may adopt distinct conformations, which we further examined by cryo-EM.

### Mutant tau filaments show different seeding activities against WT tau in cultured cells

We next examined whether the structural changes suggested by the biochemical and morphological analyses (Fig. 1) affected the ability of mutant dGAE tau filaments to seed intracellular tau aggregation. We used a previously established cell-based seeding assay, in which exogenous tau filaments are introduced into cultured cells expressing full-length tau, resulting in seed-dependent accumulation of sarkosyl-insoluble phosphorylated tau^13^. WT or mutant dGAE tau filaments were introduced into SH-SY5Y cells transiently expressing full-length 3R1N (tau with 3 microtubule-binding (MTB) repeats and 1 N-terminal insertion) or 4R1N (tau with 4 MTB repeats and 1 N-terminal insertion) WT tau, and the cells were fractionated into sarkosyl-soluble (Sar-sup) and sarkosyl-insoluble (Sar-ppt) fractions (Fig. 2A).

**Figure 2.**
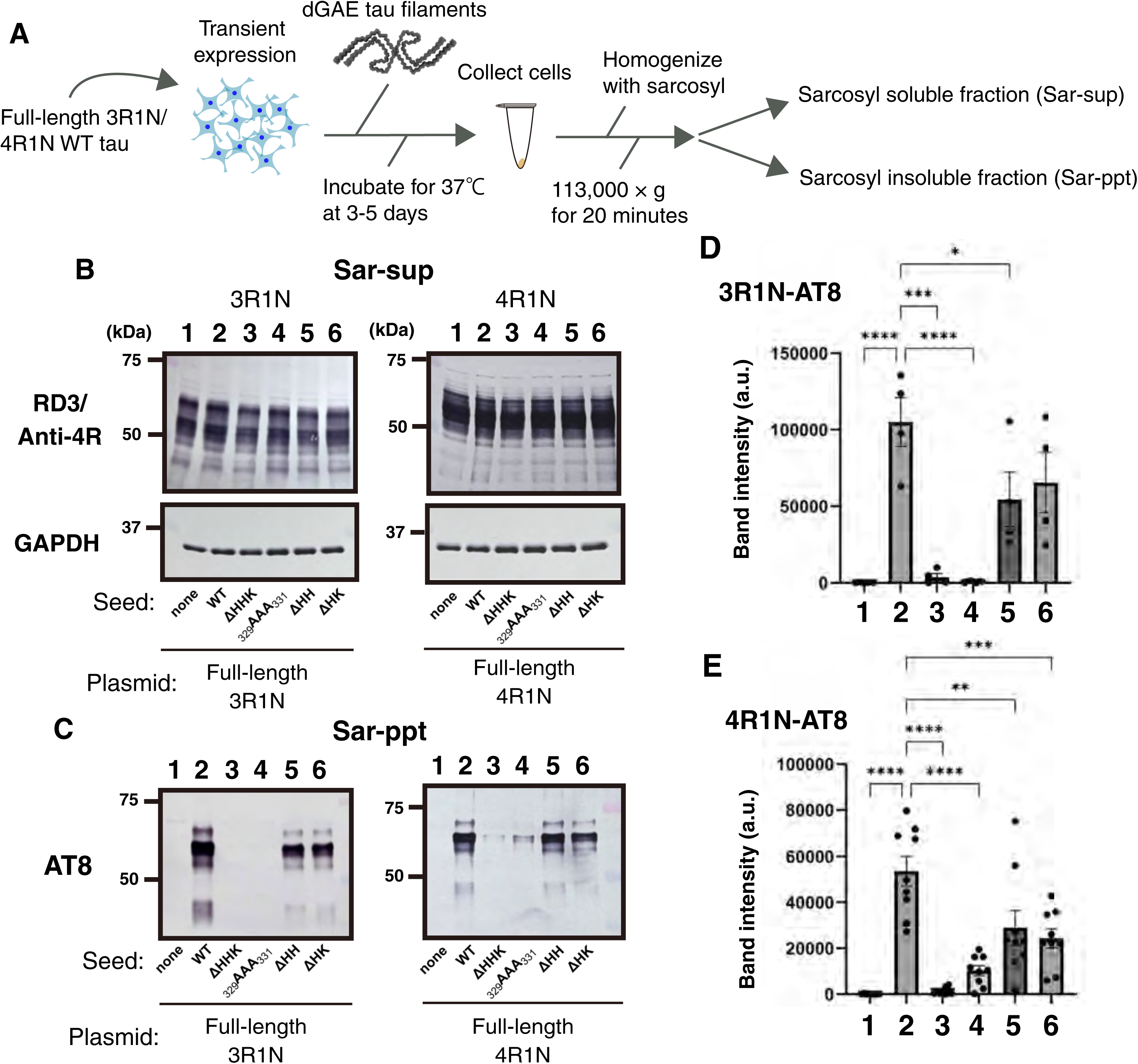
Seeding activity of WT and mutant dGAE tau filaments in cultured cells. (A) Schematic overview of the cell-based seeding assay. SH-SY5Y cells transiently expressing full-length 3R1N or 4R1N WT tau were treated with WT or mutant dGAE tau filaments. After 3–5 days, cells were harvested and fractionated into sarkosyl-soluble (Sar-sup) and sarkosyl-insoluble (Sar-ppt) fractions. (B) Immunoblot analysis of the Sar-sup fractions. Plasmid-derived soluble tau was detected using RD3 antibody for 3R tau or anti-4R antibody for 4R tau. GAPDH was used as a loading control. Lane 1, no seed; lane 2, WT; lane 3, ΔHHK; lane 4, _329_AAA_331_; lane 5, ΔHH; lane 6, ΔHK. (C) Immunoblot analysis of the Sar-ppt fractions. Insoluble phosphorylated tau was detected using the AT8 antibody. Lane order as in (B). (D, E) Quantification of AT8-positive band intensities in cells expressing 3R1N tau (D) or 4R1N tau (E). Data are presented as mean ± SEM. Each mutant was compared with WT using one-way ANOVA followed by Dunnett’s multiple comparisons test. 3R tau: N = 4; 4R tau: N = 9. *, P < 0.05; **, P < 0.01; ***, P < 0.001; ****, P < 0.0001.

Immunoblot analysis of the Sar-sup fractions showed comparable expression levels of plasmid-derived soluble tau among the samples (Fig. 2B). In the absence of seeds, little or no insoluble phosphorylated tau was detected in the Sar-ppt fraction (Fig. 2C, lane 1). Introduction of WT dGAE filaments induced robust accumulation of AT8-positive insoluble tau in cells expressing either 3R1N or 4R1N WT tau (Fig. 2C, lane 2). ΔHHK and _329_AAA_331_ filaments induced little or no AT8-positive tau accumulation (Fig. 2C, lanes 3 and 4), indicating that three-residue alteration of the _329_HHK_331_ motif severely impaired seeding activity, whereas ΔHH and ΔHK filaments produced approximately 45–60% of the accumulation seen with WT (Fig. 2C, D, E, lanes 5 and 6). These results show that the seeding activity of dGAE tau filaments depends strongly on the integrity of the _329_HHK_331_ motif. Three-residue alterations almost abolished seeding activity against full-length WT tau, whereas two-residue deletions partially preserved it, suggesting that structural features retained in the ΔHH and ΔHK filaments may remain compatible with WT tau templating.

We further asked whether the weak seeding activity of ΔHHK and _329_AAA_331_ filaments simply reflected sequence incompatibility with WT tau monomers, or whether these mutant filaments were intrinsically defective in functioning as seeds. To address this, we tested their seeding activity toward full-length tau carrying the corresponding mutations. The results revealed that ΔHHK and _329_AAA_331_ filaments did not show significantly enhanced seeding activity toward the corresponding mutant tau compared with WT tau (Supplementary Fig. 1A, B). Thus, the reduced seeding activity of these mutant filaments was not rescued by matching the mutation in the substrate tau monomer, suggesting that ΔHHK and _329_AAA_331_ filaments are intrinsically inefficient templates for seeded tau aggregation.

### Cryo-EM structures reveal distinct folds of WT and mutant dGAE tau filaments

To determine how alterations in the _329_HHK_331_ motif affect tau filament architecture, we next analyzed WT and mutant dGAE tau filaments by cryo-EM (Fig. 3). WT dGAE filaments showed several polymorphs, including straight filaments that were not suitable for helical reconstruction and three analyzable filament types composed of C-shaped protofilaments (Supplementary Fig. 2A). The major WT structure, designated WT-1, was reconstructed at 2.7 Å resolution and consisted of four C-shaped protofilaments arranged as two protofilament dimers, forming a quadruple helical filament (Fig. 3A, B and Supplementary Fig. 3A).

**Figure 3.**
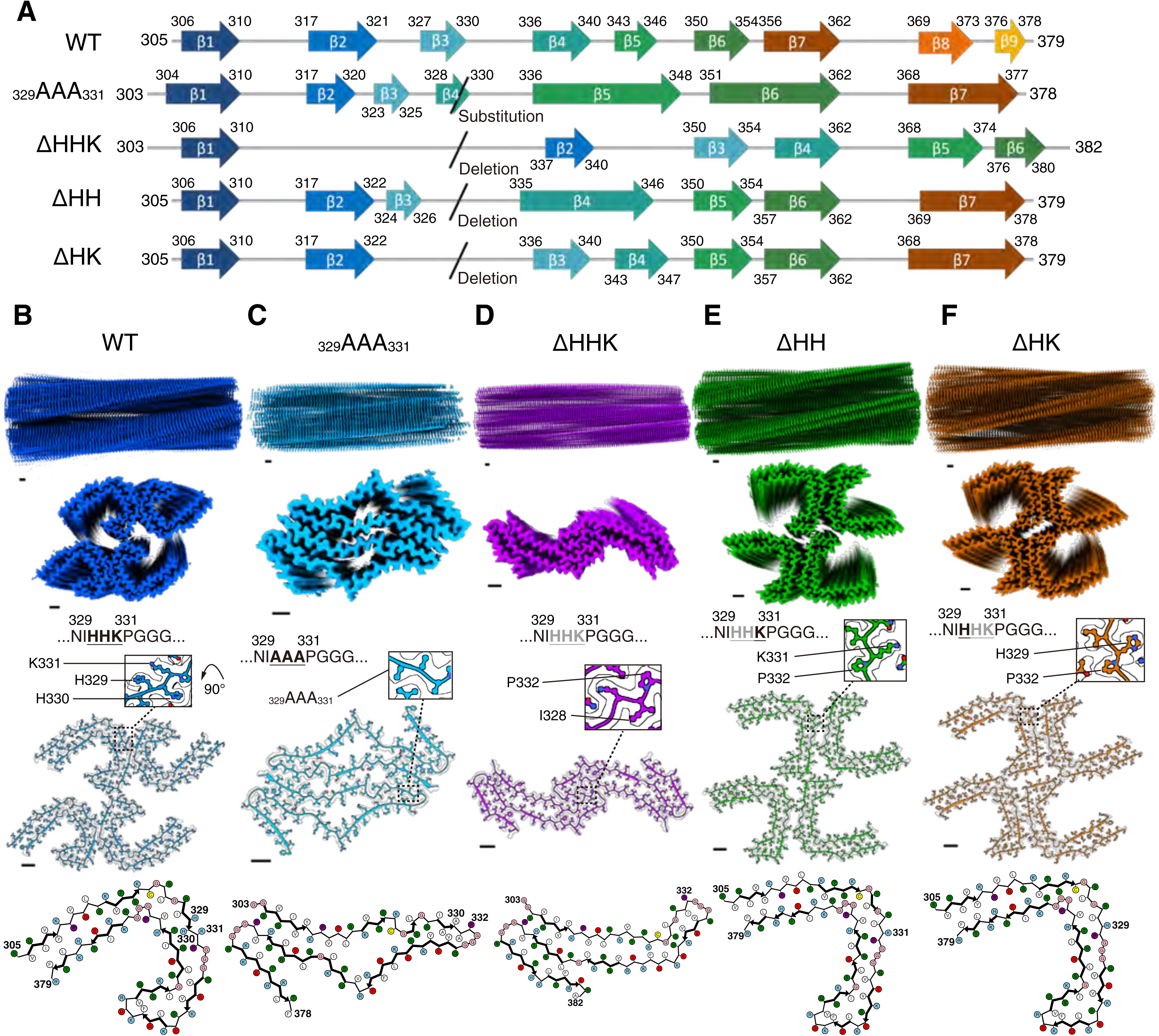
Cryo-EM structures of wild-type and mutant tau filaments. (A) Schematic summary of the β-strand positions and ordered core regions in WT, _329_AAA_331_, ΔHHK, ΔHH, and ΔHK dGAE tau filaments. Residue ranges included in the ordered core are indicated for each structure. (B–F) Cryo-EM density maps and atomic models of WT and mutant dGAE tau filaments: WT (B), _329_AAA_331_ (C), ΔHHK (D), ΔHH (E), and ΔHK (F). For each structure, side views and cross-sectional views of the density maps are shown, together with atomic models fitted into the maps, enlarged views of the mutation sites, and schematic representations of the protofilament cores. WT filaments consist of four C-shaped protofilaments arranged as two protofilament dimers, whereas _329_AAA_331_ and ΔHHK filaments form two-protofilament structures with seahorse-shaped and I-shaped protofilament folds, respectively. ΔHH and ΔHK filaments retain four-protofilament architectures composed of WT-like C-shaped protofilaments. Colors represent WT (blue), _329_AAA_331_ (cyan), ΔHHK (purple), ΔHH (green), and ΔHK (orange). In the schematic representations, residues are colored according to their physicochemical properties: positively charged, cyan; negatively charged, red; polar, green; non-polar, white; glycine, pink; proline, purple; and cysteine, yellow. Scale bar, 10 Å.

In the WT structure shown in Fig. 3B, each protofilament subunit adopts a C-shaped fold spanning residues 305–379, and within each protofilament dimer, the two subunits are related by a pseudo-2₁ screw axis and staggered by approximately half a pitch along the fibril axis, generating a zipper-like zig-zag configuration. Each protofilament subunit contains nine glycines, but only the _333_GGG_335_ triplet adopts a nearly straight main-chain conformation. The other glycines occupy bent positions that contribute to the curvature of the C-shape. This straight _333_GGG_335_ lies at the center of the protofilament dimer interface, where it engages in a short β-sheet-type interaction with the _333_GGG_335_ of its counterpart (Fig. 4A, inset a, right). Whereas the β-strands flanking _333_GGG_335_ hydrogen-bond with adjacent strands within the same protofilament, the _333_GGG_335_ triplet uniquely switches its β-sheet partner across the dimer interface from intra- to inter-protofilament. The switch is permitted by the main-chain flexibility of consecutive glycines, making _333_GGG_335_ the central structural element of each dimer. The central _333_GGG_335_ β-sheet bridge is laterally framed by interactions between the _329_HHK_331_ motif of one protofilament and the _336_QVE_338_ motif of its partner (Fig. 4A, inset a, right). Owing to the pseudo-2₁ screw symmetry and zig-zag arrangement, one _329_HHK_331_ motif engages two staggered _336_QVE_338_ motifs from adjacent protofilament subunits of the partner protofilament. Viewed along the fibril axis, each _333_GGG_335_ bridge is sandwiched between two such _329_HHK_331_-_336_QVE_338_ pairs (Fig. 4A, inset a, left). Although individually weaker than the central β-sheet interaction, these lateral _329_HHK_331_-_336_QVE_338_ contacts help position the _333_GGG_335_ bridge and frame the dimer interface. By contrast, the contact surface between the two protofilament dimers is markedly smaller, mediated only by Glu342 and Lys343 (Supplementary Fig. 3A, red dashed box). This comparatively weak interaction accounts for the modular character of the four-protofilament filament. Within each protofilament subunit, the two arms of the C-shape are brought together by intramolecular packing of the N-terminal PHF6-containing region (residues 305–316) against the C-terminal region (residues 370–379). This region is stabilized by hydrophobic interactions among Val306, Ile308, Tyr310, Leu376, and Phe378, and by hydrogen bonds involving Tyr310–His374 and Asp314–Ser316/Lys370 (Fig. 4A, inset b).

**Figure 4.**
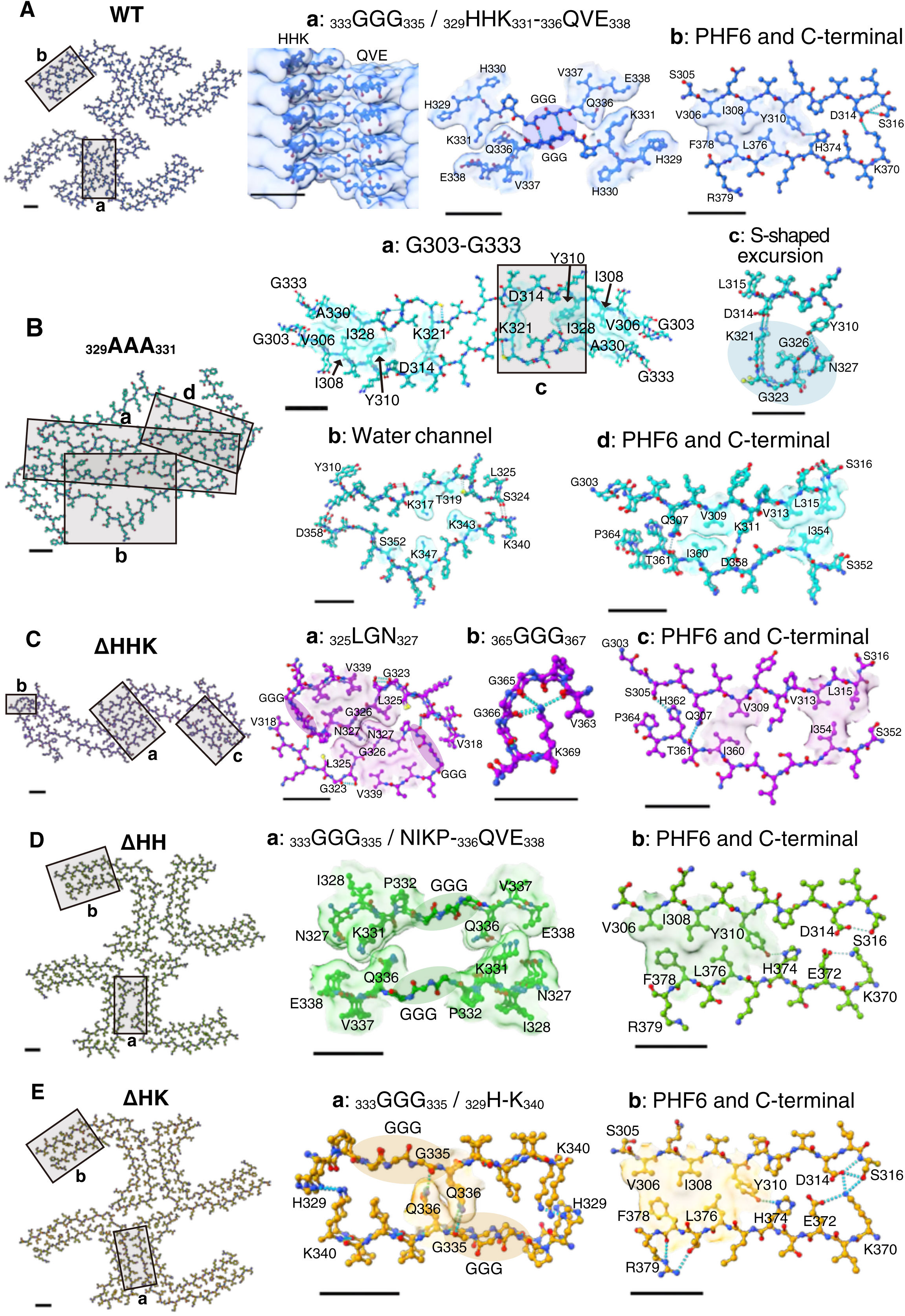
Protofilament interfaces and intramolecular core packing of WT and mutant dGAE tau filaments. Atomic models and cryo-EM density maps showing the protofilament interfaces and core-packing regions of (A) WT, (B) _329_AAA_331_, (C) ΔHHK, (D) ΔHH, and (E) ΔHK dGAE tau filaments. Boxed regions in the overview at the left of each panel are enlarged in the lettered insets. Scale bar, 10 Å. (A) WT. (a) Protofilament dimer interface centered on the _333_GGG_335_ β-sheet-type bridge, laterally flanked by _329_HHK_331_-_336_QVE_338_ contacts, with H329 and E338 indicated. (b) Intramolecular packing of the PHF6-containing N-terminal region, represented by S305 and S316, against the C-terminal region, represented by K370 and R379, which closes the C-shaped core. (A) _329_AAA_331_. (a) Inter-protofilament interface spanning residues G303-G333. (b) Solvent-accessible water channel accommodating residues including K317, T319, K343, K347 and S352. (c) S-shaped main-chain excursion involving G323 and G326 (semitransparent circle). (d) Repacking of the PHF6-containing N-terminal region with downstream core segments, represented by G303, S316, S352, and P364 within the seahorse-shaped core. (B) ΔHHK. (a) Inter-protofilament interface in which the V318-V339 segments interlock through a _325_LGN_327_ steric zipper. (b) The _365_GGG_367_ turn, stabilized by K369 hydrogen-bonding to main-chain carbonyls around V363. (c) Repacking of the PHF6-containing N-terminal region with downstream core segments, represented by G303, S316, S352, and P364 within the I-shaped core. (C) ΔHH. (a) Protofilament dimer interface in which the two _333_GGG_335_ triplets face each other without direct contact. The dimer is bridged by the flanking N327-I328-K331-P332 (NIKP) and _336_QVE_338_ regions, with N327 and E338 indicated. (b) PHF6-containing N-terminal and C-terminal packing, represented by S305, S316, K370, and R379, maintaining the WT-like C-shaped core. (D) ΔHK. (a) Protofilament dimer interface in which the shifted _333_GGG_335_ triplets bring H329 and K340 of the opposing protofilaments into proximity, with the double-layered structure stabilized by G335–Q336 hydrogen bonds and a steric zipper between opposing Q336 residues. (b) PHF6-containing N-terminal and C-terminal packing, represented by S305, S316, K370, and R379, preserving the C-shaped core. In all panels, cyan dashed lines indicate hydrogen bonds, semitransparent surfaces shown in each filament color indicate non-hydrogen-bonded interaction regions (such as hydrophobic interactions, electrostatic interactions, water-mediated interactions, interactions between specific motifs and steric zipper structure), and black boxes highlight the key local structural features described in each inset.

Cryo-EM analyses of _329_AAA_331_-mutant filaments revealed four polymorphs (Supplementary Fig. 2B). The major structure (_329_AAA_331_-1), determined at 2.9 Å, formed a left-handed helix with pseudo-2₁ screw symmetry and is composed of two nearly identical protofilaments (Fig. 3C, Supplementary Fig. 3B). Each protofilament subunit adopts a “seahorse”-shaped fold, a novel conformation distinct from the WT C-shaped fold. The interface between the two protofilaments is formed by residues G303–G333 and stabilized by an ionic interaction between D314 of one protofilament and K321 of the other, together with hydrophobic interactions involving V306, I308, Y310, I328 and A330 (Fig. 4B, inset a). Lys317, Thr319, Lys343, Lys347 and Ser352 are accommodated within a solvent-accessible water channel that connects them to bulk solvent (Fig. 4B, inset b). Neither _333_GGG_335_ nor _365_GGG_367_ participates in inter-protofilament contacts, and both occupy loop regions (Supplementary Fig. 3B, black box). Two isolated glycines (Gly323 and Gly326) lie at turn positions that introduce an S-shaped excursion of the main chain, allowing Lys321–Asp314, Tyr310–Gly326 and Ser324–Lys340 to form an alternative hydrogen-bonding network that stabilizes the interior (Fig. 4B, inset b and c). The replacement of _329_HHK_331_ with three alanines abolishes the lateral _329_HHK_331_-_336_QVE_338_ interaction that, in WT, flanks the _333_GGG_335_ β-sheet bridge, removing the structural context required for assembly of the WT protofilament dimer. In contrast to WT, the interactions between the N-terminal PHF6-containing region and the C-terminal region are substantially rearranged. In this region, the alternative packing is stabilized by hydrophobic interactions involving V313–L315–I354 and V309–I360, together with hydrogen bonds between Q307 and T361, and between K311 and D358 (Fig. 4B, inset d).

We observed two polymorphs in ΔHHK-mutant filaments (Supplementary Fig. 2C). The major structure (ΔHHK-1) was determined at 2.8 Å and formed a left-handed helix with pseudo-2₁ screw symmetry (Fig. 3D, Supplementary Fig. 3C). Like _329_AAA_331_ and unlike WT, the filament comprises only two protofilaments. At the interface between the two protofilaments, the V318-V339 segments of the opposing protofilament subunits are arranged in opposite orientations and interlock through their L325-N327 residues, whose side chains fit into the gaps between one another, forming a steric zipper (Fig. 4C, inset a). Each protofilament subunit adopts an “I”-shaped fold with a double-layered β-sheet kinked at Gly323 and Lys343 and an interlayer spacing narrower than in WT and _329_AAA_331_ (Supplementary Fig. 3C, black dashed box). This compact packing is a direct consequence of the three-residue _329_HHK_331_ deletion. Because the side chains of a β-strand project alternately toward the inside and outside of the layered β-sheet, the three-residue deletion inverts the inside/outside register of the β-strand. The resulting register shift allows all interior lysine residues to find appropriate polar partners within the layered core, producing the tightly packed double-layered fold seen here. Both glycine triplets occupy turn regions of the protofilament subunit, as in _329_AAA_331_, and neither participates in inter-protofilament contacts (Supplementary Fig. 3C, black box). The _333_GGG_335_ turn is stabilized by hydrophobic interactions between its flanking residues, while the _365_GGG_367_ turn is supported by Lys369, whose side chain hydrogen-bonds with the main-chain carbonyl oxygens of Val363, Gly365, and Gly366 (Fig. 4C, inset b). The three-residue deletion abolishes the _329_HHK_331_-_336_QVE_338_ interaction at the protofilament dimer interface, leading to the displacement of _333_GGG_335_ into a turn, which eliminates the WT protofilament dimer interface and results in the formation of an I-shaped protofilament structure distinct from the AD-like C-shaped fold. In addition, the three-residue deletion induces an inversion of the side-chain register, which is also considered to contribute to the formation of a structure distinct from the C-shaped fold. Consistent with the loss of the _329_HHK_331_-_336_QVE_338_ interaction and inversion of the side-chain register, the N-terminal PHF6-containing region and the C-terminal segment no longer close around the conserved buried hydrogen-bond network seen in WT. Instead, these regions are repacked within the I-shaped core, where they form new contacts that stabilize this alternative fold. These contacts comprise hydrophobic interactions among V313-L315-I354/V309-I360 and hydrogen bonds between S305–H362 and Q307–T361 (Fig. 4C, inset c).

### Two-residue deletion mutants retain WT-like C-shaped protofilament architecture

We next examined the structures of the two-residue deletion mutants, ΔHH and ΔHK, which retained partial seeding activity in cultured cells. Cryo-EM analysis of ΔHH-mutant filaments revealed a single ordered polymorph (ΔHH-1) besides unanalyzable straight filaments (Supplementary Fig. 2D). The ΔHH-1 structure, determined at 2.4 Å, formed a left-handed helix with C1 symmetry. In contrast to the three-residue mutants, ΔHH filaments are composed of four protofilaments arranged as two protofilament dimers (Fig. 3E, Supplementary Fig. 3D). Within each protofilament dimer, the two _333_GGG_335_ triplets face each other but, unlike WT, make no direct contact because the inter-strand distance is now too large. The protofilament dimer is instead bridged by the flanking N327–I328–K331–P332 (NIKP) and _336_QVE_338_ motifs, centered on Asn327 and Glu338 (Fig. 4D, inset a), whose spacing holds the triplets apart. The two protofilament dimers then associate through two inter-dimer contacts, the WT-like Glu342–Lys343 pair and an additional Lys321–Asp348 pair (Supplementary Fig. 3D, red dashed box). In contrast, the two protofilament dimers in WT associate through only a single contact (Supplementary Fig. 3A, red dashed box). Each protofilament subunit adopts a C-shaped, double-layered β-sheet fold nearly identical to that of WT, though slightly altered in curvature to a more L-shaped appearance. In ΔHH filaments, the surviving K331 and surrounding residues interact with the opposing _336_QVE_338_ motif, thereby partially compensating for the lost inter-protofilament interactions and allowing retention of the WT-like fold (Fig. 4D, inset a). Furthermore, this retention of the WT-like fold is associated with the even-residue nature of the deletion. Because the side chains of a β-strand project alternately toward the inside and outside of the layered β-sheet, a two-residue deletion preserves their inside/outside register, whereas the three-residue deletion in ΔHHK inverts it. The two-residue deletion in ΔHH produced only a main-chain register shift and slightly opened the C-shaped structure. The retention of inter-protofilament interactions and preservation of the side-chain register also maintain WT-like interactions between the N-terminal PHF6-containing region and the C-terminal region, thereby preserving a WT-like core (Fig. 4D, inset b).

Cryo-EM analysis of ΔHK-mutant filaments determined a major structure (ΔHK-1) at 2.2 Å that formed a left-handed helix with C1 symmetry and comprised four protofilaments arranged as two protofilament dimers (Fig. 3F). Each protofilament subunit adopts a C-shaped, double-layered β-sheet fold nearly identical to WT but slightly more open (L-shaped). Although the overall architecture closely resembles ΔHH, the two _333_GGG_335_ triplets within each protofilament dimer are oriented differently. Rather than facing each other, they are shifted so that each triplet interacts with the three residues following its partner’s triplet, bringing His329 and Lys340 of the opposing protofilaments into proximity (Fig. 4E, inset a). Furthermore, the double-layered structure is stabilized by hydrogen bonds between Gly335 and Gln336, together with a steric zipper formed between opposing Gln336 residues (Fig. 4E, inset a). This shift also reshapes the inter-dimer interface. Glu342 and Lys343, which directly link the two protofilament dimers in WT, no longer interact directly but instead engage the adjacent Asp345 and Lys347 (Supplementary Fig. 3E, red dashed box), giving narrower water-accessible cavities than in ΔHH. In ΔHK filaments, the WT-like fold is also preserved, as observed in ΔHH, because the surviving H329 interacts with the opposing K340, partially compensating for the lost inter-protofilament interactions and allowing retention of the WT-like fold (Fig. 4E, inset a). As in ΔHH, this two-residue deletion preserves the inside/outside register of the β-strand side chains and produces only a main-chain register shift. ΔHK retains the interactions between the N-terminal PHF6-containing region and the C-terminal region observed in WT and ΔHH, thereby maintaining a WT-like core (Fig. 4E, inset b).

ΔHK-mutant filaments were markedly polymorphic, unlike ΔHH, showing five polymorphs besides a straight filament (Supplementary Fig. 2E). In addition to the major ΔHK-1, these comprised a triple-helical filament of three C-shaped protofilaments (ΔHK-2), a novel pairing of one C-shaped protofilament with a differently folded protofilament (ΔHK-3), and a PHF-like assembly of two protofilaments (ΔHK-4). ΔHK-5 could not be reconstructed due to an insufficient number of particles.

### The _329_HHK_331_ region governs the C-shaped protofilament fold and seeding activity

To examine how changes in the _329_HHK_331_ motif affect the C-shaped fold, we superimposed WT and mutant protofilaments on their N-terminal (PHF6-containing) regions using ChimeraX and assessed how the downstream core was repositioned relative to this conserved segment. In the three-residue mutants _329_AAA_331_ and ΔHHK, the N-terminal segment (residues 305–328) retained a WT-like conformation but was flipped around the _329_HHK_331_ region, displacing the downstream protofilament core. This rearrangement accounts for their non-C-shaped, two-protofilament folds (Fig. 5A, B). _329_AAA_331_ showed a pronounced deviation with an RMSD of 3.59 Å for residues S305–I328, although the proximal S305–S320 segment retained partial similarity to WT (RMSD = 1.82 Å). By contrast, ΔHHK deviated less from WT, with the PHF6-containing N-terminal segment retaining partial similarity (RMSD = 1.44 Å for residues S305–I328).

**Figure 5.**
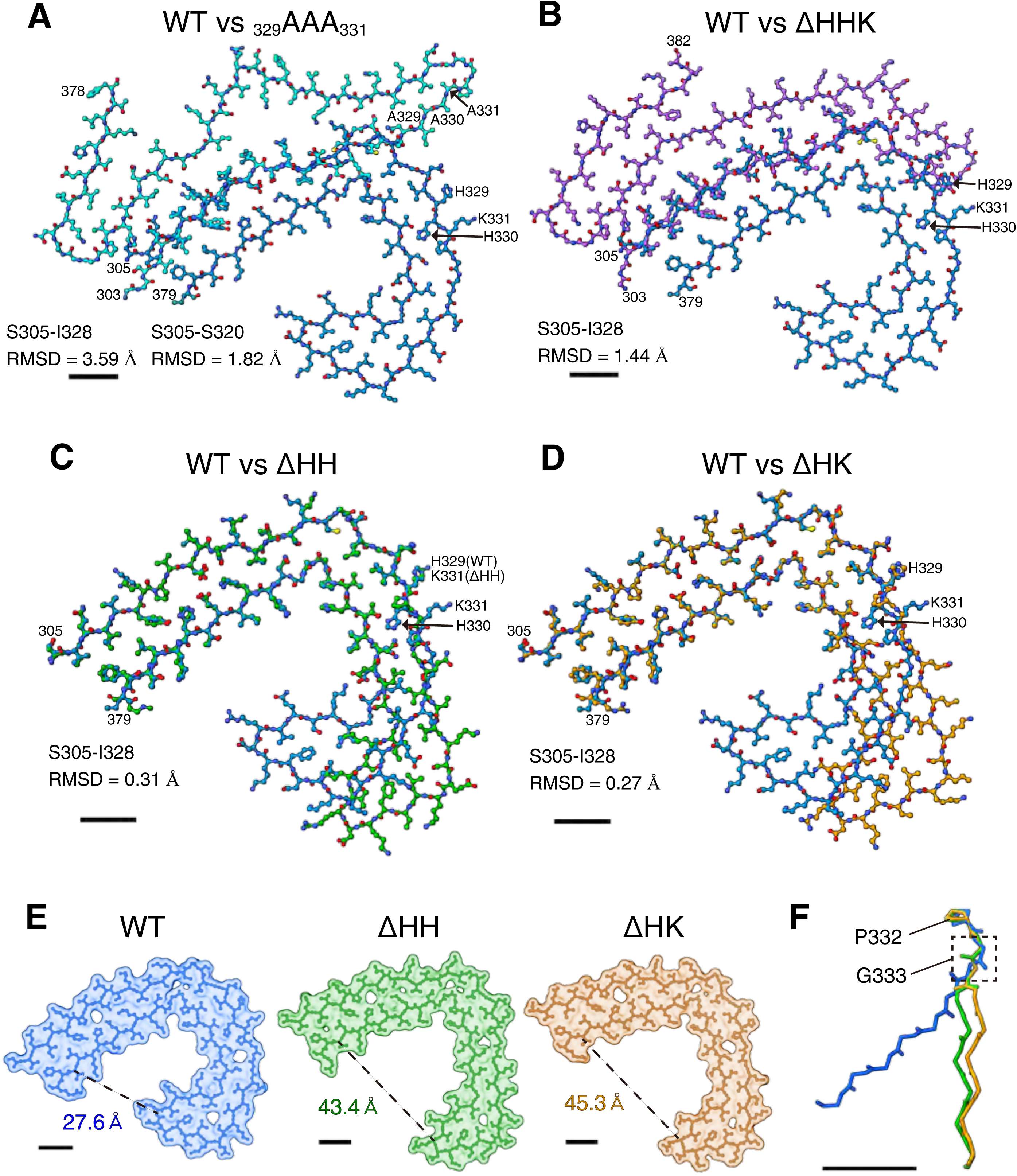
Structural superposition of WT and mutant dGAE tau protofilaments. (A–D) Superposition of the N-terminal region of WT (blue) protofilaments with each mutant: (A) _329_AAA_331_ (cyan), (B) ΔHHK (purple), (C) ΔHH (green), and (D) ΔHK (orange). Root-mean-square deviation (RMSD) values for the N-terminal segment (residues S305-I328) are indicated. For _329_AAA_331_, the RMSD for the proximal S305-S320 segment is also shown. Residues at positions 329-331 are labelled in each comparison. (E) Surface representations of the protofilament cores of WT (blue), ΔHH (green), and ΔHK (orange). Dashed lines indicate the R349-T377 distance, which reflects the degree of opening of the C-shape (27.6 Å for WT, 43.4 Å for ΔHH, and 45.3 Å for ΔHK). (F) Backbone (Cα) overlay of WT (blue), ΔHH (green), and ΔHK (orange) protofilaments. The dashed box highlights the shift in the main-chain dihedral angles of Gly333 within the _332_PGGG_335_ motif. Scale bars, 10 Å.

In contrast to these three-residue mutants, the corresponding N-terminal regions of the two-residue deletion mutants ΔHH and ΔHK closely overlapped with WT, with RMSD values of 0.31 Å and 0.27 Å, respectively, for residues S305–I328 (Fig. 5C, D). Consistent with this, both mutants retained the four-protofilament, C-shaped architecture, with the surviving _329_HHK_331_ residue contributing to the protofilament dimer interface in place of the WT _333_GGG_335_ bridge. Their C-shaped protofilaments were slightly more open than those of WT, as reflected by increased R349–T377 distances in ΔHH and ΔHK (Fig. 5E). This opening corresponded to a shift in the dihedral angles of Gly333 within the _332_PGGG_335_ motif, while the overall WT-like fold was preserved (Fig. 5F).

These results indicate that alteration of all three residues in the _329_HHK_331_ motif causes loss of the WT-like inter-protofilament interactions. This loss changes the direction of folding around the N-terminal PHF6-containing region, thereby inducing structures distinct from the C-shaped fold. In ΔHHK, the deletion additionally inverts the side-chain inside/outside register, further driving this structural change. In contrast, in the two-residue deletion mutants, the remaining single residue (His329 or Lys331) partially compensates for the lost inter-protofilament interactions, allowing the C-shaped architecture to be maintained in a slightly relaxed form, while the side-chain inside/outside register is preserved. Thus, the cellular seeding activity of tau filaments corresponds to the extent to which the C-shaped protofilament fold is preserved. Specifically, _329_AAA_331_ and ΔHHK filaments, which do not maintain a WT-like fold, lose seeding activity, whereas ΔHH and ΔHK filaments, which preserve a WT-like C-shaped fold, retain partial seeding activity.

### The _347_KDR_349_ motif is not required for the formation of the C-shaped dGAE tau protofilament fold

To determine whether the effect of _329_HHK_331_ mutation was specific to this region or reflected a more general effect of altering charged residues within the tau repeat domain, we next examined the _347_KDR_349_ motif. This sequence has been proposed as an aggregation-prone region^11^, but unlike _329_HHK_331_, it is not located at the central protofilament dimer interface of the WT dGAE filament. We therefore generated a _347_AAA_349_ mutant, in which the _347_KDR_349_ residues were substituted with alanines, and analyzed the resulting filaments by cryo-EM.

The _347_AAA_349_ mutant formed several polymorphs, with _347_AAA_349_-1 representing the major population (Fig. 6A). The structure was reconstructed at 2.3 Å resolution, as assessed by Fourier shell correlation analysis (Fig. 6B). Cryo-EM density maps and atomic modeling revealed that the _347_AAA_349_ filament was composed of two protofilaments, each adopting a C-shaped fold closely resembling that of WT dGAE protofilaments (Fig. 6C, D). The substituted residues 347–349 were located in a turn of the C-shaped protofilament fold (Fig. 6E) and did not induce the marked fold rearrangement observed in the _329_AAA_331_ mutant (Fig. 3C).

**Figure 6.**
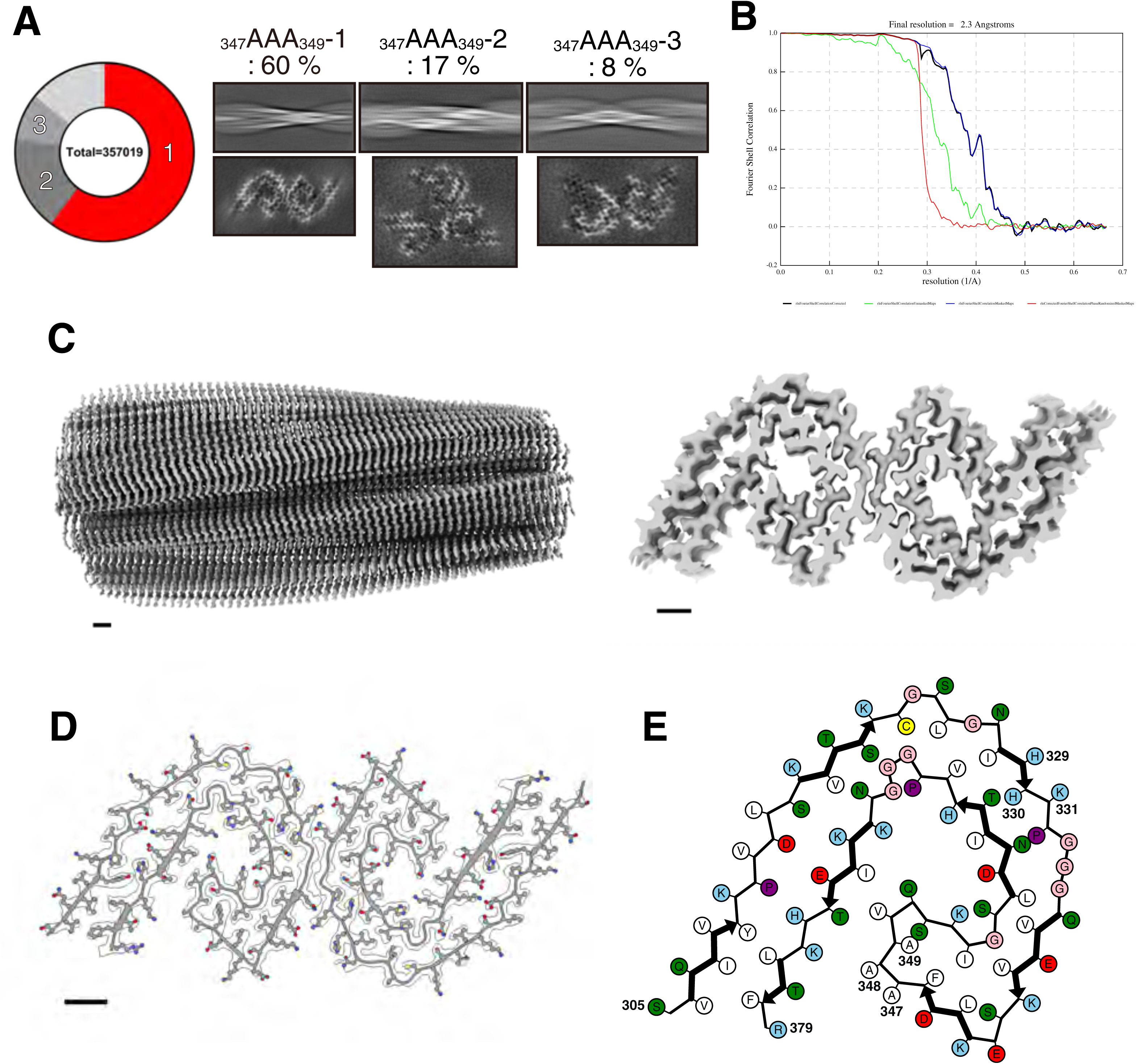
Cryo-EM structure of the _347_AAA_349_ mutant dGAE tau filament. (A) Distribution of observed polymorphs and representative cryo-EM data for the _347_AAA_349_ mutant. The pie chart shows the proportion of each filament type relative to the total number of particles contributing to the 2D class averages (total = 357,019). Unnumbered light-gray sectors represent straight filaments and false positives. For each type, a representative 2D class average is shown at the top, and, where reconstructed, a cross-sectional view of the density map is shown at the bottom. (B) Fourier shell correlation (FSC) curves. The final resolution of the _347_AAA_349_-1 reconstruction was 2.3 Å. (C) Cryo-EM density map of the _347_AAA_349_-1 filament, composed of two C-shaped protofilaments: side view (left) and cross-section (right). (D) Atomic model fitted into the cryo-EM density map. (E) Schematic representation of the protofilament core, with residues 305, 347–349, and 379 labelled. Residues are colored according to their physicochemical properties: positively charged, cyan; negatively charged, red; polar, green; non-polar, white; glycine, pink; proline, purple; and cysteine, yellow. Scale bar, 10 Å.

These results indicate that alanine substitution of the _347_KDR_349_ motif does not substantially disrupt the C-shaped dGAE tau protofilament fold. Thus, among the charged motifs examined here, the _329_HHK_331_ region has a more specific role in maintaining the AD-like C-shaped protofilament conformation.

## Discussion

Cryo-EM structures of tau filaments from AD brains have revealed that residues _329_HHK_331_ are positioned near an inter-protofilament interface of PHFs^5^. Similar clusters of basic residues are also found at or near inter-protofilament interfaces in disease-associated α-synuclein filaments, including Lys43 and Lys45 in multiple system atrophy filaments and Lys32, Lys34, Lys43, and Lys45 in Lewy body disease filaments^14, 15^. These observations led us to consider that clustered basic residues may play an important structural role in the assembly of disease-associated amyloid filaments.

In tau, the _329_HHK_331_ motif is located immediately N-terminal to the _332_PGGG_335_ sequence, a glycine-rich segment positioned near the protofilament interface in mature AD-type tau filaments^10^. This motif has also been implicated in binding to negatively charged aggregation-promoting factors such as heparin^12^. We therefore hypothesized that the _329_HHK_331_ motif may not merely promote tau aggregation, but may instead act as a local structural determinant of AD-like tau filament folding. In this study, we tested this hypothesis by analyzing the aggregation propensity, cellular seeding activity, and cryo-EM structures of recombinant dGAE tau filaments carrying deletions or alanine substitutions in this motif.

ThT fluorescence is widely used to monitor amyloid fibril formation because its fluorescence increases upon binding to β-sheet-rich amyloid assemblies^16, 17^. Previous studies have demonstrated that structurally distinct tau polymorphs can exhibit markedly different ThT reactivity. For instance, Fitzpatrick et al.^5^ and Falcon et al.^3, 4^ showed that tau filaments from different tauopathies differ not only in their cryo-EM structures but also in their biochemical and biophysical properties. More recently, Lövestam et al.^8^ showed that recombinant dGAE tau filaments can recapitulate disease-specific folds, validating dGAE as a model system in which sequence changes can be linked to defined structural outcomes. The divergent ThT profiles and filament morphologies we observed for the _329_HHK_331_ mutants are therefore consistent with a structural basis for altered protofilament folding, rather than a simple reduction in aggregation efficiency (Fig. 1).

In the WT dGAE filament, the _333_GGG_335_ motif forms a central inter-protofilament bridge within each protofilament dimer, while the adjacent _329_HHK_331_ motif supports the surrounding interface through contacts with the _336_QVE_338_ region (Figs. 3 and 4). Thus, the _329_HHK_331_-_336_QVE_338_ interaction appears to provide the local structural context that allows the _333_GGG_335_ segment to adopt its straight conformation and participate in dimer formation. Alteration of all three residues in the _329_HHK_331_ motif, either by deletion or alanine substitution, disrupted the _329_HHK_331_-_336_QVE_338_ interaction at the protofilament dimer interface, leading to the displacement of _333_GGG_335_ into a turn, thereby converting the filaments into non C-shaped two-protofilament structures (Figs. 3 and 4). In the deletion mutant ΔHHK, this was accompanied by inversion of the side-chain inside/outside register. ΔHHK and _329_AAA_331_ adopted different folds, I-shaped and seahorse-shaped, respectively. Both lost the WT-like C-shaped protofilament architecture and showed little seeding activity toward WT tau (Fig. 2). This low seeding activity was not rescued by using tau monomers carrying the corresponding mutations (Supplementary Fig. 1), suggesting that these mutant filaments are intrinsically poor templates rather than simply mismatched with WT tau. These findings suggest that disruption of the _329_HHK_331_-supported interface compromises the structural features required for efficient tau templating.

By contrast, the two-residue deletion mutants ΔHH and ΔHK retained WT-like C-shaped protofilament folds and four-protofilament architectures despite local rearrangements at the protofilament dimer interface. In these mutants, two structural features act together. First, although the _333_GGG_335_ bridge seen in WT is not formed, the surviving residue (Lys331 in ΔHH or His329 in ΔHK) and its neighbors establish alternative contacts that maintain the protofilament dimer interface. Second, because the deletion removes an even number of residues, the inside/outside register of the β-strand side chains is preserved, unlike the inversion caused by the three-residue deletion. This register preservation maintains the WT-like packing between the N-terminal PHF6-containing region and the C-terminal region, thereby stabilizing the C-shaped fold. Together, these features allow the C-shaped fold to persist in a more open or relaxed form. The partial preservation of this WT-like architecture is consistent with the partial seeding activity of these filaments, suggesting that compatibility with WT tau templating depends on the extent to which the C-shaped protofilament fold is retained.

Several sequence-based and peptide-based studies have identified aggregation-prone regions within the tau repeat domain, including PHF6, PAM4, _326_GNIHHK_331_, and the _344_LDFKDR_349_ region ^9, 11^. However, aggregation propensity at the peptide level does not necessarily mean that the same sequence determines the disease-associated fold of the intact tau filament. The comparison between _329_HHK_331_ and _347_KDR_349_ in this study supports this distinction (Fig. 6). Whereas both regions contain charged residues and have been implicated in tau aggregation, only alteration of the _329_HHK_331_ motif markedly disrupted the AD-like C-shaped protofilament fold. These findings suggest that the _329_HHK_331_ region acts as a local fold-determining element, rather than merely as one of several aggregation-prone sequences within tau.

This study also has several limitations. Because we used recombinant dGAE tau, this system may not fully recapitulate tau filament formation in the AD brain. Therefore, further investigation will be required to determine whether similar results are obtained in full-length tau systems capable of reproducing AD-type structures such as recombinant phosphomimetic tau PAD12^18^. Also in the brains of patients with AD, post-translational modifications (e.g., phosphorylation), cofactors, molecular chaperones, and the local intracellular environment may influence filament formation and seeding activity, but these factors were not fully addressed in this study. Thus, our findings should be interpreted with these considerations in mind.

Despite these limitations, our findings provide insight into the mechanisms underlying tau filament formation in AD, identifying the _329_HHK_331_ region as a local sequence element that links AD-like tau filament architecture with seeding activity. Further studies using disease-derived tau filaments and *in vivo* models will be required to determine whether this region can serve as a structural target for interfering with AD-type tau filament elongation or spread.

## Methods

### Construction of wild-type and mutant dGAE tau plasmids

WT tau dGAE fragment (residues 297–391, based on the amino acid numbering of human tau 4R2N), ΔHHK mutant (His329, His330 and Lys331 deletion), _329_AAA_331_ mutant (_329_HHK_331_ residues substituted by Ala), _347_AAA_349_ mutant (_347_KDR_349_ residues substituted by Ala), ΔHH mutant (His329 and His330 deletion), and ΔHK mutant (His330 and Lys331 deletion) were amplified by PCR using the following primers and subcloned into pET22b: WT dGAE, forward, 5′-ATGATCAAACACGTCCCGGGAGGCG-3′ and reverse, 5′- TTACTCCGCCCCGTGGTCTGTCTTG-3′; ΔHHK, forward, 5′- CCGGGTGGCGGGCAGGTGGAGGTTAAAAGC-3′ and reverse, 5′- AATATTGCCCAAACTGCCACATTTGGAGGT-3′; _329_AAA_331_, forward, 5′- GCCGCCGCACCGGGTGGCGGGCAGGTGGAG-3′ and reverse, 5′- AATATTGCCCAAACTGCCACATTTGGAGGT-3′; _347_AAA_349_, forward, 5′- GCAGCTGCCGTGCAAAGCAAGATTGGCTCG-3′ and reverse, 5′- AAAGTCCAGTTTCTCGCTTTTAACCTCCAC-3′; ΔHH, forward, 5′- AAACCGGGTGGCGGGCAGGTGGAGGTTAAA-3′ and reverse, 5′- AATATTGCCCAAACTGCCACATTTGGAGGT-3′; ΔHK, forward, 5′- CCGGGTGGCGGGCAGGTGGAGGTTAAAAGC-3′ and reverse, 5′- GTGAATATTGCCCAAACTGCCACATTTGGA-3′. All constructs were verified by DNA sequencing.

### Expression and purification of recombinant dGAE tau monomers

Plasmids encoding WT or mutant dGAE tau were transformed into *E. coli* BL21 (DE3). Transformed cells were plated on LB agar containing 50 µg/mL Na-carbenicillin and incubated overnight at 37°C. The obtained colonies were collected and inoculated into 500 mL Plusgrow II medium (Nacalai Tesque) supplemented with Na-carbenicillin at a final concentration of 50 µg/mL. After 2 h of incubation at 37°C, protein expression was induced by adding isopropyl-β-D-thiogalactopyranoside to a final concentration of 0.8 mM, followed by incubation for 5 h at 37°C or overnight at 24°C. The culture was then transferred to centrifuge tubes and centrifuged at 2,700 g for 10 min at 4°C. For protein purification, frozen bacterial cell pellets were thawed and resuspended in 10 mL of Buffer A (50 mM PIPES-NaOH, pH 6.9, 1 mM EDTA, 1 mM dithiothreitol (DTT), and 0.5 mM PMSF). The suspension was sonicated for 2 min on ice before centrifugation at 26,600 g for 15 min at 4°C. Then, the supernatant was collected and 2-mercaptoethanol at a 1/100 ratio was added. The mixture was heated at 100°C for 5 min. After heating, the sample was centrifuged again at 26,600 g for 15 min at 4°C and the supernatant collected. This supernatant was applied to an SP Sepharose™ Fast Flow column (Cytiva; column volume: 3 mL) equilibrated with 30 mL of Buffer A. Then, the column was washed with 30 mL of Buffer A, and bound proteins were eluted with 9 mL of Buffer A containing 0.35 M NaCl. To precipitate tau proteins, ammonium sulfate was added to 50% saturation, and the mixture was incubated on ice for 1 h. After centrifugation at 26,600 g for 20 min at 4°C, the pellet was dissolved in 2 mL of Buffer B (10 mM phosphate buffer (PB), pH 7.4, 10 mM DTT) and loaded onto a HiLoad Superdex gel filtration column on the size exclusion chromatography system (ÄKTA go, Cytiva). The resulting fractions were concentrated using a centrifugal concentrator (3 kDa cutoff) to ∼15–20 mg/mL; this fraction was used as a recombinant dGAE tau monomer. The protein concentration was measured by reverse-phase high-performance liquid chromatography (RP-HPLC) using an Aquapore RP- 300 column (3 mm diameter × 4 cm length; Brownlee) on the HPLC system (Agilent Technologies). Recombinant dGAE tau monomers were stored at −80°C until further use.

### Preparation of recombinant dGAE tau filaments

Recombinant dGAE tau filaments were prepared by shaking a recombinant dGAE tau monomer solution (2 mg/mL) in the presence of 10 mM PB, 10 mM DTT, and 200 mM magnesium chloride (MgCl₂)^8^. Specifically, 100 µL of the reaction mixture was added per well in a 96-well plate and fibrillization was induced by shaking at 37°C and 200 rpm for 48 h using a FLUOstar Omega plate reader (BMG LABTECH). To monitor changes in fibrillization over time, a reaction mixture supplemented with ThT (final concentration at 4 µM) was prepared in separate wells and ThT fluorescence intensity was measured once every 30 min. After 48 h, the reaction mixture without ThT was collected, and an aliquot was used for cryo-EM analyses. The remaining reaction mixture was centrifuged at 135,000 g for 20 min at room temperature, and dGAE tau filaments were collected as a pellet. To remove residual monomers, 500 µL of saline (Otsuka Pharmaceutical) was added to the pellet and centrifugation was repeated under the same conditions. Subsequently, the pellet was resuspended in the same volume of saline as before centrifugation and subjected to sonication. This mixture was used as seed in subsequent analyses. To measure the protein concentration, 10 µL aliquot of the mixture was mixed with 89 µL of 6 M guanidine hydrochloride and 1 µL of 0.1 M DTT, and analyzed using RP-HPLC.

### Quantification of recombinant dGAE tau filaments

Each recombinant dGAE tau monomer was fibrillized using a FLUOstar Omega plate reader as described above, and a 50 µL aliquot of the reaction mixture collected and centrifuged at 135,000 g for 20 min at room temperature. To remove residual monomers, the resulting pellet was washed by addition of 500 µL saline followed by centrifugation under the same conditions. Then, the pellet was resuspended in 50 µL of 10 mM PB before adding 10 µL of 5× SDS sample buffer. Samples were sonicated and subsequently heated at 100°C for 5 min. From each sample, 1 µL was loaded onto a 15% polyacrylamide gel and subjected to SDS-PAGE and proteins detected by Coomassie brilliant blue staining. Band intensities were quantified using ImageJ and data plotted using Prism 10.6.1 (GraphPad).

### Negative-stain electron microscopy analysis of dGAE tau filaments

A 3 µL aliquot of each dGAE tau filament sample was applied to a collodion-coated copper grid (400 mesh; Nisshin EM). After the sample had spread across the grid, excess was removed using a pipette, and any remaining moisture blotted with a paper towel. Then, 4 µL of a 2% sodium phosphotungstate solution was applied to the grid and incubated at room temperature for 15 s. The excess stain was again blotted off with a paper towel and the staining procedure repeated once. The grid was then left to dry at room temperature for several min before examination using a JEM-1400 transmission electron microscope. The half-pitch length and width of the filaments were measured using ImageJ and data plotted using Prism 10.6.1.

### Analysis of dGAE tau filament seeding activity in cultured cells

The seeding activity of dGAE tau filaments using cultured cells was tested as previously described^13^. Human neuroblastoma SH-SY5Y cells were cultured in DMEM/F-12 (Sigma-Aldrich) medium supplemented with 10% (v/v) fetal bovine serum (FBS), a non-essential amino acid solution (GIBCO), and a penicillin-streptomycin-glutamine solution (GIBCO). A total of 8 × 10⁵ cells were seeded per well in a 6-well plate and incubated overnight. The next day, a full-length tau expression plasmid (pcDNA3-human 4R1N/3R1N tau WT) was transfected using X-tremeGENE 9 (Roche). After 3∼24 h, 2 µg of dGAE tau filaments were introduced into the cells using MultiFectam reagent (Promega). Treated cells were incubated in a CO₂ incubator for 3–5 days before being collected and centrifuged at 1,800 g for 5 min at 25°C. The cell pellet was then resuspended in 300 µL of A68 buffer (10 mM Tris-HCl, pH 7.5, 10% sucrose, 0.8 M NaCl, 1 mM EGTA) containing 1% sarkosyl, and subjected to sonication for ∼20 s to lyse the cells. Subsequently, an additional 300 µL of A68 buffer containing 1% sarkosyl was added to the lysate, before centrifugation at 113,000 g for 20 min at 25°C. The resulting supernatant (300 µL, Sar-sup) was collected and mixed with 75 µL of a 5× SDS sample buffer containing 5% 2-mercaptoethanol, followed by heating at 100°C for 5 min. For the pellet fraction (Sar-ppt), 500 µL PBS was carefully added without disturbing the pellet, followed by centrifugation at 113,000 g for 20 min at 25°C to eliminate any remaining supernatant. The pellet was then resuspended in 50 µL of 2× SDS sample buffer containing 5% 2-mercaptoethanol, sonicated, and heated at 100°C for 5 min. Both samples (Sar-sup and Sar-ppt) were separated by electrophoresis using a 7.5% polyacrylamide gel and transferred onto polyvinylidene difluoride membranes (Millipore) at 200 mA for 1 h. Then, membranes were blocked with PBS containing 3% gelatin (Wako) for 10 min and incubated with primary antibodies diluted in PBS containing 10% bovine serum overnight. The primary antibodies used were Anti-4R-tau (TIP-4RT-P01, Cosmo Bio, 1:2000 dilution), RD3 (8E6/C11, Millipore, 1:2000 dilution), AT8 (Invitrogen, 1:1000 dilution) and GAPDH (Millipore, 1:1000 dilution). After washing with Tris-buffered saline (TS), the membranes were incubated for 2 h with a biotinylated secondary antibody (Biotin-Goat anti-mouse IgG, Biotin-Goat anti-rabbit IgG, Vector) diluted in PBS containing 10% FBS (1:500 dilution), followed by another wash with TS. Then, membranes were incubated for 1 h with an avidin-biotin enzyme complex (ABC, Vector), washed with PBS, and developed using a solution containing 0.04 mg/mL 3,3′-diaminobenzidine (DAB, Sigma-Aldrich), 1.6 mg/mL nickel chloride and 0.03% hydrogen peroxide in PBS. The reaction was stopped by washing the membranes with water. Band intensities were quantified using ImageJ and data was plotted using Prism 10.6.1.

### Cryo-EM data acquisition

For sample preparation, tau monomers were added to the dGAE tau filament solution to a final concentration of 50 µM and the mixture incubated at room temperature for 10 min. The monomer was added to prevent filament clumping as previously reported^19^. For cryo-EM data acquisition, 3 µL of dGAE tau filament solution (1–2 mg/mL) was applied to glow-discharged carbon grids (Quantifoil Cu or Au R1.2/1.3, 300 mesh). The grids were then blotted with filter paper and plunge-frozen in liquid ethane using a Vitrobot Mark IV (Thermo Fisher Scientific) at 100% humidity and 18°C. Movies were recorded using a Titan Krios G4 cryo-electron microscope (Thermo Fisher Scientific) equipped with a Falcon 4i detector at pixel sizes of 0.75 or 0.96 Å/pixel and a defocus range of −0.8 to −2.0 µm. Detailed conditions are shown in Table 2.

**Table 2.**
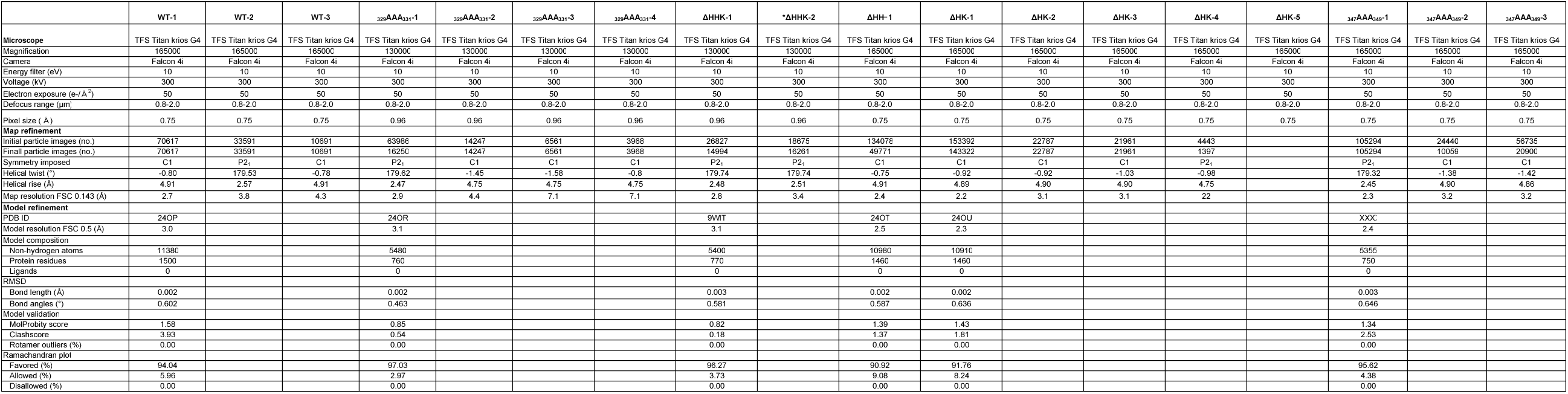
Cryo-EM data acquisition, processing, map refinement, and model validation statistics. *ΔHHK-Type2 was analyzed by merging two datasets.

### Cryo-EM data processing

Each movie was processed using RELION’s motion correction job for gain correction, alignment, and dose-weighting^20^. To exclude micrographs with excessive sample movement, images showing motion >80 Å were removed using a subset selection job. Contrast transfer function parameters were estimated using CTFFIND4.1^21^. Based on CtfMaxResolution values, images with a resolution >4–6 Å were excluded, as were images with defocus values >2.5 µm. Helical reconstruction was performed using RELION version 5.0.0. Filaments were manually or automatically picked using a Topaz-trained model^22^. Particles were initially extracted with a box size of either 1024 or 768 pixels and then downscaled to 256 or 128 pixels, respectively. The first 2D classification included 100–150 classes, iterating until the resolution plateaued to estimate the crossover distance and distinguish polymorphs. Based on 2D class averages, polymorphs were separated, and particles corresponding to each polymorph independently classified. For datasets using auto-picking, particles were re-extracted using smaller box sizes, and 2D classification was repeated to remove as many false positives and low-quality class averages as possible, thereby facilitating subsequent initial model generation. Initial models were generated using the relion_helix_inimodel2d program^23^. Then, 3D auto-refinements were performed to optimize the helical parameters. If no improvement in resolution or separation of β-strands was observed, a 3D classification was performed at T = 8–20 and K = 1. To further improve the resolution, two rounds of Bayesian polishing and CTF refinement were conducted^20, 24^. Non-alignment 3D classification was then performed and the best class selected. In the final round of 3D auto-refinement, final maps were sharpened using a postprocessing job and the reported resolutions estimated using a threshold of 0.143 in the Fourier shell correlation between two independently refined half-maps^25^. Detailed processing parameters are summarized in Supplementary Fig. 4 and Table 2.

### Detection and quantification of polymorphs

Polymorphs were separated using FilamentTools in the Filament tab of the Subset selection job in RELION version 5.0.0^10^. This procedure was performed using either the initial 2D classification or the 2D classification performed after removing false positives. We calculated the reported percentages of polymorphs by extracting particle counts for each filament from the logfile.pdf, run.out, and run_optimiser.star files generated by this job and expressing each count as a percentage of the total count. This percentage does not necessarily reflect the actual number of particles used for the final 3D reconstruction. Further details are provided in Supplementary Fig. 2.

### Atomic modeling

Atomic models were built automatically using the ModelAngelo job in RELION version 5.0.0^26^. Atomic models were refined as previously described^27^. Specifically, one protofilament subunit was cropped in ChimeraX, and initial refinement performed using COOT and Phenix^28, 29, 30^. The refined model was then opened in ChimeraX and a one-subunit model was generated using the “Fit in Map.” This one-subunit model was further refined in COOT and Phenix. Then, the refined one-subunit model was opened in ChimeraX and used to build a five-subunit model, refined in Phenix to produce the final atomic model. Note that atoms showing no clear density were deleted using COOT. Structural representations and schematics were prepared using ChimeraX and atom2svg.py.

### Data availability

Cryo-EM density maps and refined models have been deposited in the Electron Microscopy Data Bank and RCSB Protein Data Bank (RCSB PDB) under the following accession numbers: dGAE WT filament, EMD-69727 and 24OP; ΔHHK, EMD-66009 and 9WIT; _329_AAA_331_, EMD-69728 and 24OR; ΔHH, EMD-69729 and 24OT; ΔHK, EMD-69730 and 24OU; Accession codes for the _347_AAA_349_ structure will be provided upon publication.

## Supporting information

This article contains supporting information.

## Acknowledgments

We thank the members of the Dementia Research Project in Tokyo Metropolitan Institute of Medical Science for their helpful discussions and support. We are grateful to Shunsuke Kanno, Shohei Takaki, Reiko Ohtani and Taeko Kimura for their technical assistance. We used generative AI tools (ChatGPT, Gemini, and Claude) for language editing and text revision.

## CRediT authorship contribution statement

**Yuta Sato**: Writing–original draft. Methodology. Investigation. Formal analysis. Data curation. **Masato Kawasaki**: Investigation. Methodology. **Toshio Moriya**: Investigation. Methodology. Formal analysis. Data curation. **Miki Senda**: Formal analysis. Methodology. Data curation. **Masami Masuda-Suzukake**: Methodology. Resources. **Kanae Ando**: Supervision. **Shin-ichi Hisanaga**: Supervision. **Masato Hasegawa**: Supervision. Funding acquisition. **Toshiya Senda**: Writing–original draft. Supervision. Funding acquisition. **Takashi Nonaka**: Conceptualization. Writing–original draft. Supervision. Resources. Funding acquisition. Methodology. Investigation.

