## Supplementary material for "The _329_HHK_331_ motif is essential for Alzheimer’s disease tau filament fold": Supple Fig. 1-4

**Supplementary information**

**Sato Yuta et al.**

**Supplementary Figure 1. Cross-seeding activity of WT and mutant dGAE tau filaments in cultured cells.**

(A, B) Cross-seeding assays between WT and ΔHHK-mutant tau (A) or between WT and _329_AAA_331_-mutant tau (B). SH-SY5Y cells transiently expressing full-length 4R1N WT tau or the corresponding mutant tau were seeded with dGAE tau filaments derived from WT or the same mutant. Sarkosyl-soluble (Sar-sup) and sarkosyl-insoluble (Sar-ppt) tau were detected using anti-4R and AT8 antibodies, respectively, with GAPDH as a loading control. Lanes 1–3, cells expressing full-length WT tau; lanes 4–6, cells expressing full-length mutant tau. Within each set, cells were treated with no filaments (lanes 1 and 4), WT dGAE filaments (lanes 2 and 5), or mutant dGAE filaments (lanes 3 and 6). Right, quantification of AT8-positive insoluble tau band intensities. Data are presented as mean ± SEM (N = 4). Comparisons were made using one-way ANOVA followed by Šídák's multiple comparisons test. **, P < 0.01; ***, P < 0.001; ****, P < 0.0001; ns, not significant.

**Supplementary Figure 2. Polymorphism of WT and mutant dGAE tau filaments.**

(A–E) Polymorph distribution and representative cryo-EM data for WT (A), _329_AAA_331_ (B), ΔHHK (C), ΔHH (D), and ΔHK (E) dGAE tau filaments. For each filament type, representative 2D class averages and, where available, cross-sectional views of the reconstructed density maps are shown. Pie charts indicate the relative abundance of each identified polymorph, calculated from the number of particles contributing to the 2D class averages. Unnumbered light-gray sectors represent straight filaments, false positives, or classes that were not suitable for helical reconstruction. Type numbers correspond to the polymorphs described in the Results and Table 2.

**Supplementary Figure 3. Schematic representation of amino acid residues in WT and mutant dGAE tau filament cores.**

(A–E) Schematic diagrams showing the arrangement of amino acid residues within the protofilament core regions of WT (A), _329_AAA_331_ (B), ΔHHK (C), ΔHH (D), and ΔHK (E) dGAE tau filaments. The diagrams correspond to the atomic models shown in Fig. 3 and summarize the residue-level organization of each fold. Black boxes indicate the positions of the two GGG triplets in each tau filament. Black dashed boxes highlight the residues located at the kink positions in the ΔHHK filament structure (Gly323 and Lys343), and red dashed boxes highlight the inter-dimer interaction sites between the two dimers that constitute the WT, ΔHH, and ΔHK filament structures. Residues are colored according to their physicochemical properties: positively charged, cyan; negatively charged, red; polar, green; non-polar, white; glycine, pink; proline, purple; and cysteine, yellow.

**Supplementary Figure 4. Fourier shell correlation curves of WT and mutant dGAE tau filament reconstructions.**

Fourier shell correlation (FSC) curves for cryo-EM reconstructions of WT and mutant dGAE tau filament polymorphs. For each reconstruction, corrected, unmasked, masked, and phase-randomized masked FSC curves are shown. Final map resolutions were estimated using the FSC 0.143 criterion and ranged from 2.2 to 22 Å.
