## Supplementary material for "The _329_HHK_331_ motif is essential for Alzheimer’s disease tau filament fold": Supple Fig. 1-4

### Supplementary figure 1 Sato et al.

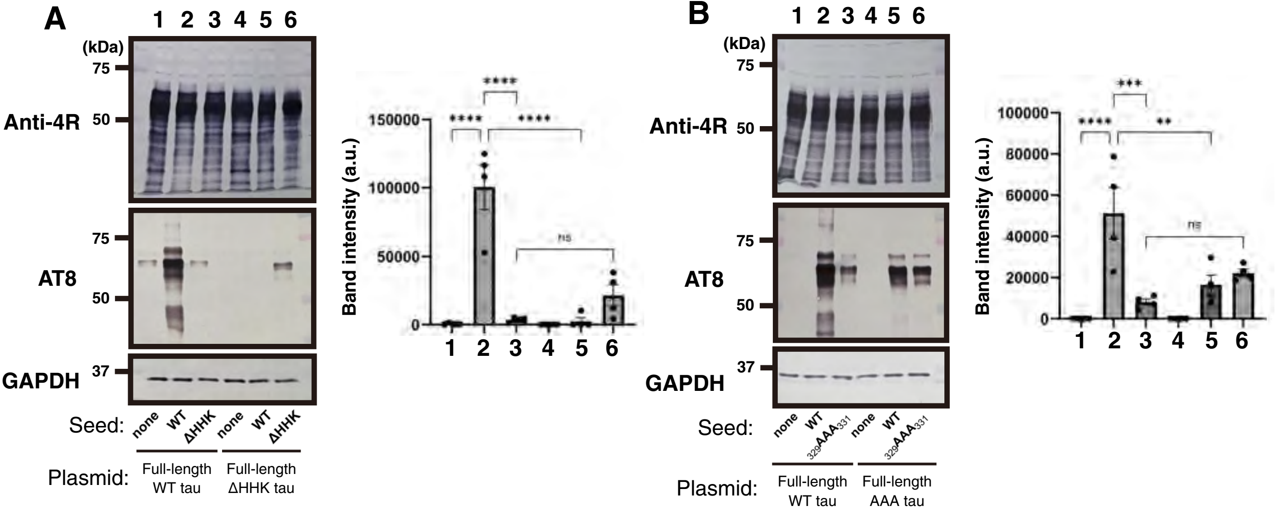

### Supplementary figure 2 Sato et al.

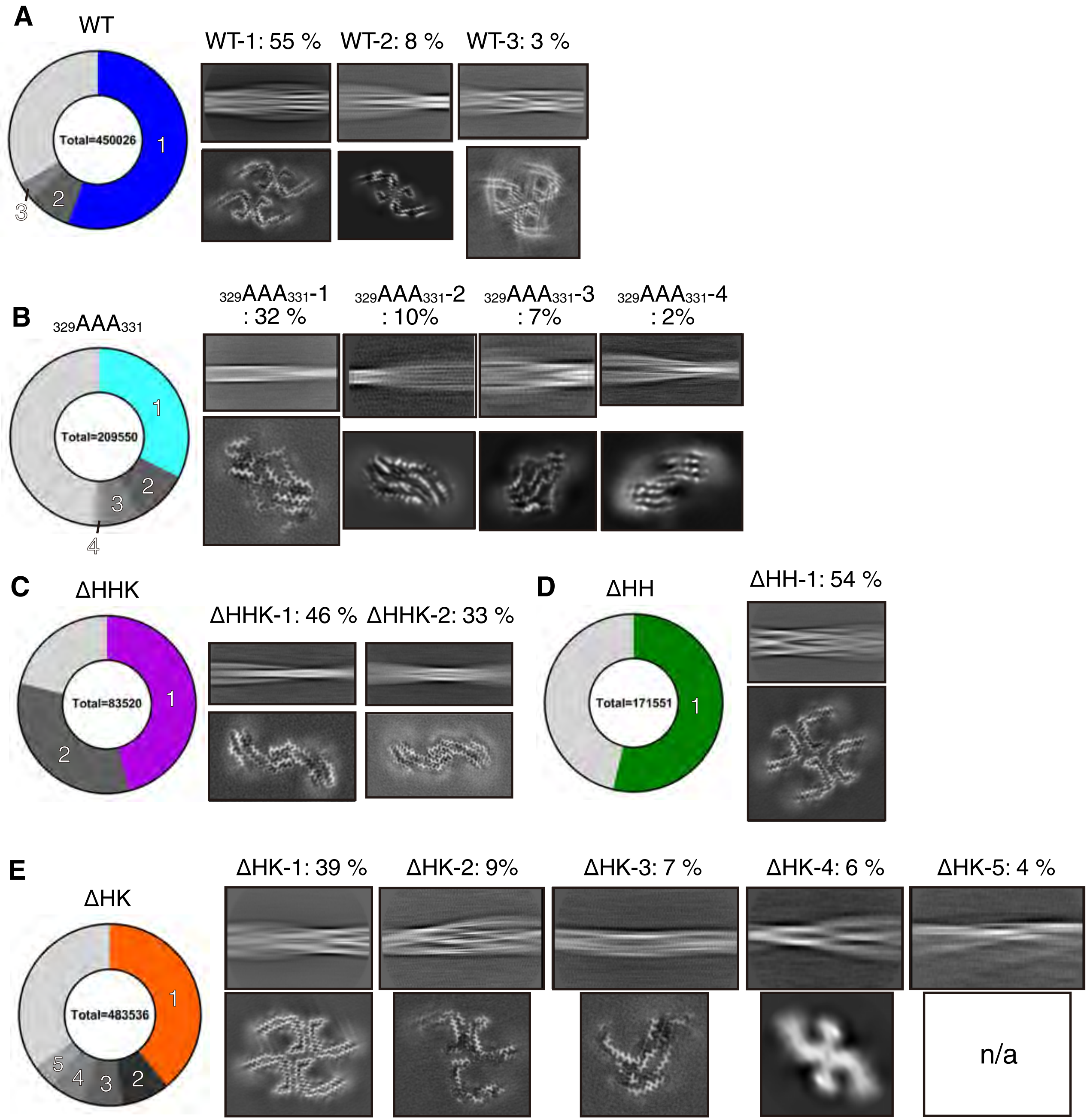

### Supplementary figure 3 Sato et al.

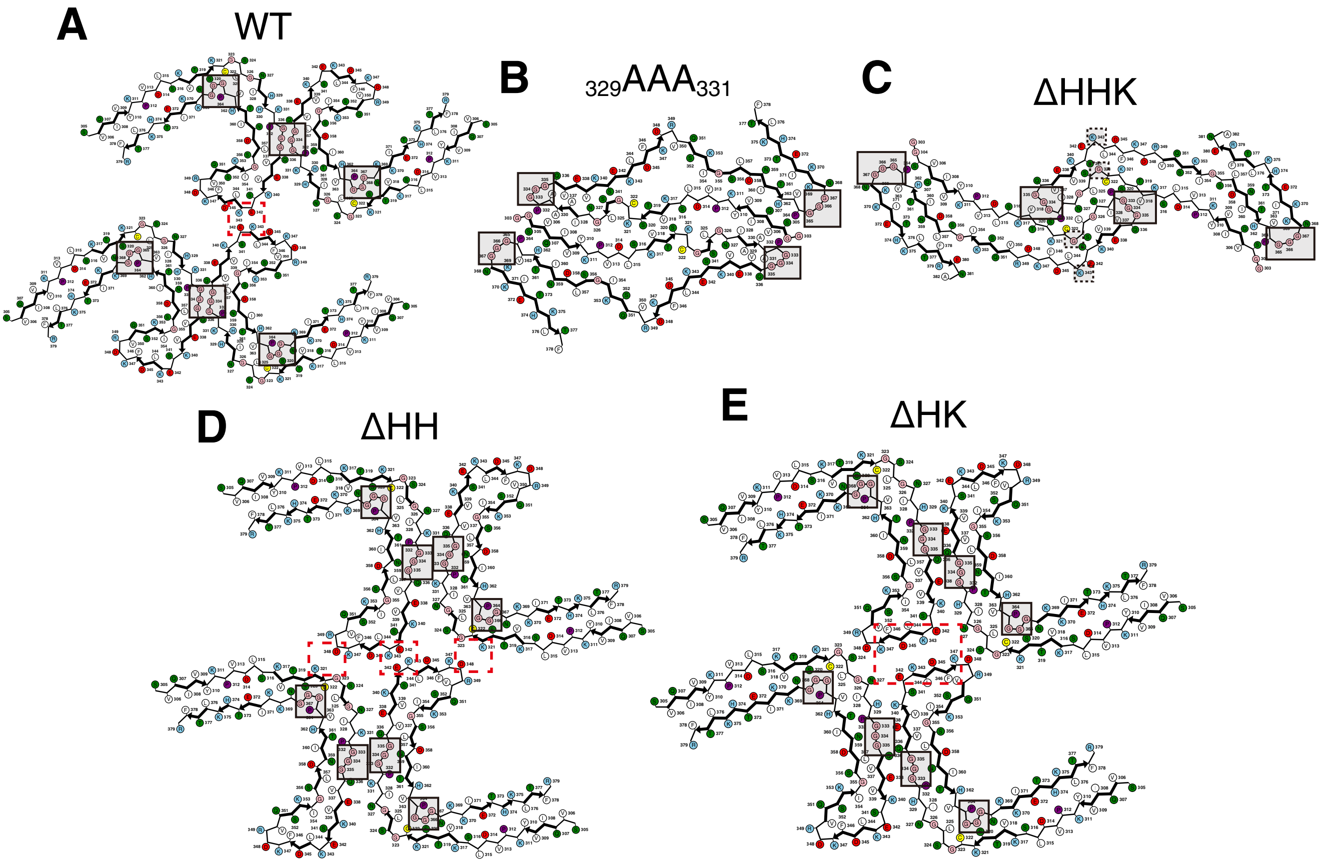

### Supplementary figure 4 Sato et al.

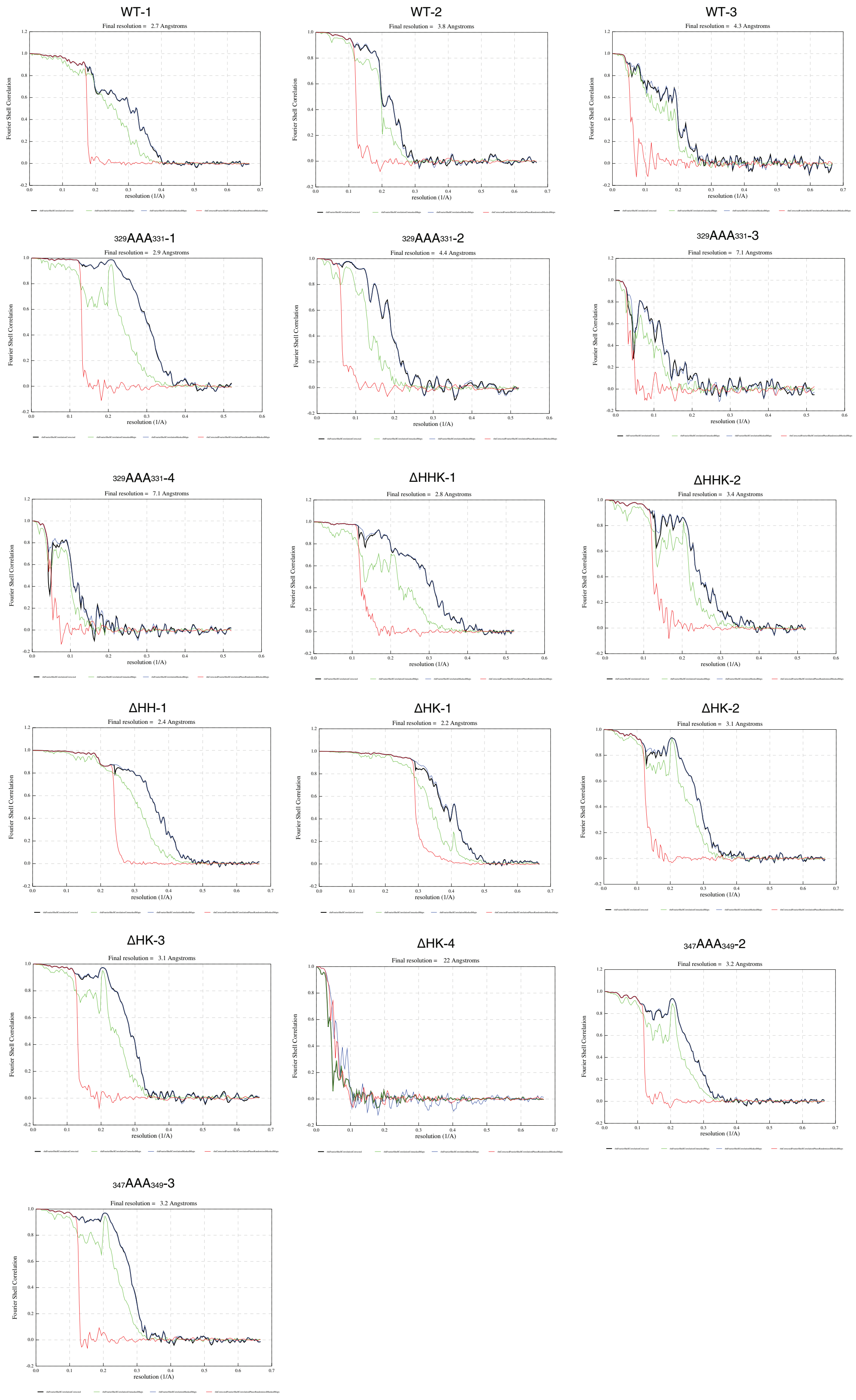
